# Overcoming effects of heterogeneous binding on BLI analysis

**DOI:** 10.1101/2025.04.25.650696

**Authors:** Noah Sherer, Jae-Hyun Cho

**Affiliations:** Department of Biochemistry and Biophysics, Texas A&M University, College Station, TX 77843, United States

## Abstract

Binding characteristics, such as k_on_, k_off_, and K_D_, are critical for mechanistic study of biomolecular interactions and drug design. Biolayer interferometry (BLI) has become popular due to its simplicity and sensitivity. Despite its widespread use, BLI data analysis is susceptible to various non-ideal features in sensorgrams. One commonly observed issue is a persistent signal drift after the binding process reaches an expected steady state. The basis of this phenomenon, often referred to as heterogeneous binding, remains poorly understood. In this study, we find that analyte aggregation on the biosensor, particularly induced by ligand-analyte complexes, can contribute to heterogeneous binding. We also find that heterogeneous binding affects not only the association phase but also the dissociation process, leading to erroneous binding characteristics. We propose an approach to mitigate the adverse impacts of heterogeneous binding on the BLI analysis. Since accurate binding characterization is fundamental for many biophysical analyses, addressing this issue is crucial.

## Introduction

Biolayer interferometry (BLI) has become a popular approach for the characterization of biomolecular interactions, primarily because it can provide both kinetic and thermodynamic parameters of biomolecular interactions with a smaller amount of samples compared to other approaches.^1-3^ As such, BLI has wide applicability from drug discovery, to the study of protein-protein interactions.^4-9^ However, BLI data often exhibits non-ideal behavior, which can hinder accurate data analysis and lead to erroneous binding parameters.^10^

One common non-ideal behavior is signal drift which results in an inclined rather than a flat steady-state signal (**Fig 1A**).^11-13^ Signal drift suggests secondary, often non-specific, binding of multiple analytes to the ligand or biosensors. The underlying mechanisms of heterogeneous binding remain poorly understood, yet they are critical for designing appropriate BLI experiments and analyzing the data. For instance, if the observed heterogeneity corresponds to functional multimeric analyte-ligand interactions, a longer association period may be required. In contrast, if it arises from an artifact, efforts should be made to minimize it for accurate data interpretation.

**Figure 1:**
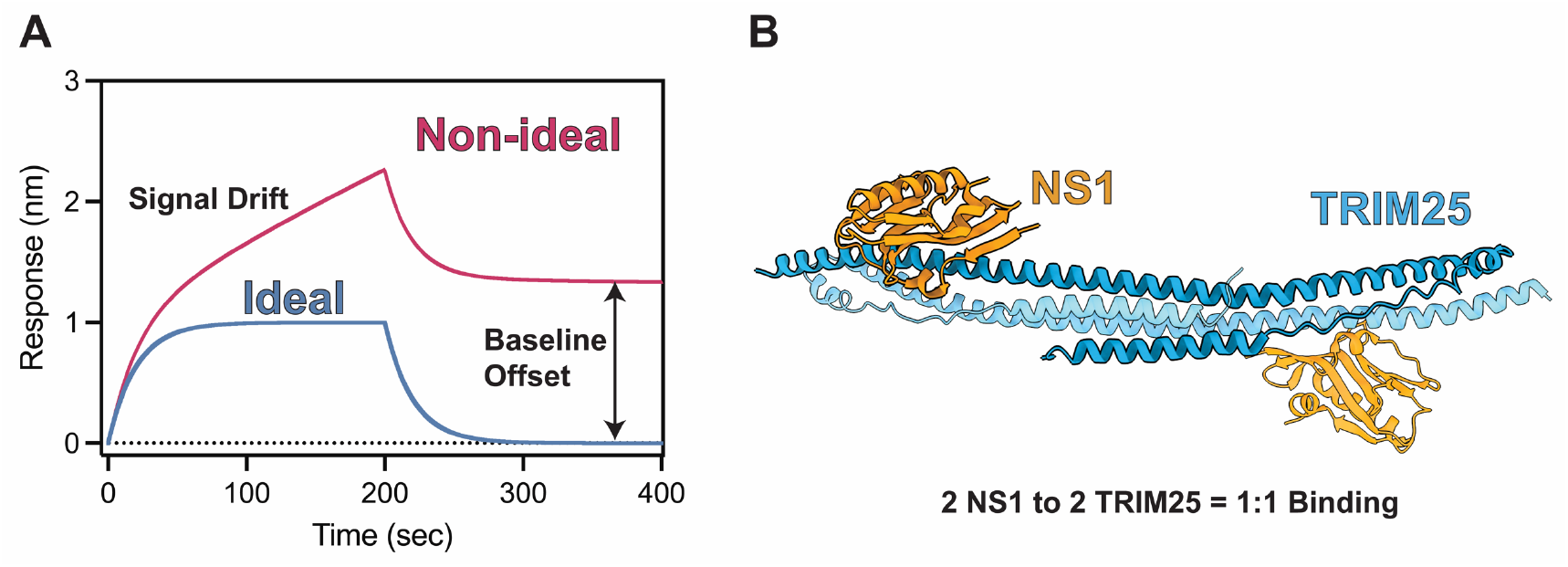
Protein-protein interactions monitored by BLI. **(A)** A representative BLI sensorgram depicting ideal and non-ideal binding behavior. Ideal binding behavior (blue) is characterized by an association phase that reaches a flat steady-state and a dissociation phase that returns to the baseline. Non-ideal behavior (magenta) is characterized by constant signal drift in the association phase and an incomplete dissociation phase **(B)** Crystal structure of the TRIM25-NS1 complex (PDB: 5NT1). One TRIM25-NS1 interaction pair is highlighted to underscore the 1:1 binding.

In practice, a multi-phasic exponential function is often used to fit data exhibiting heterogeneous binding behavior.^14-17^ However, the lack of mechanistic understanding makes it difficult to rationally justify the use of multi-phasic fitting equations. Moreover, it remains unclear how heterogeneous binding affects data analysis even when multi-phasic models are applied. As a result, the accuracy of the parameters obtained from the multi-phasic fit—such as association rate constant (k_on_), dissociation rate constant (k_off_), and equilibrium dissociation constant (K_D_)—needs to be verified. To this end, we sought to identify potential mechanisms underlying heterogeneous binding and to examine its effects on data analysis.

In this study, we investigate the interaction between the nonstructural protein 1 (NS1) of influenza A virus and human tripartite motif-containing protein 25 (TRIM25) using BLI (Fig 1B). NS1 is a major virulence factor of influenza A viruses, playing a key role in suppressing interferon expression and apoptosis in host cells.^18-21^ TRIM25 is an E3 ubiquitin ligase that regulates viral RNA sensing and innate immune responses to infection.^22-25^ The ability of NS1 to hijack TRIM25 contributes to viral evasion of host immunity.^26, 27^ Moreover, the TRIM25 binding surface on NS1 is shared with another host protein, phosphoinositide 3-kinase (PI3K).^10, 27-31^ Understanding the molecular mechanisms by which NS1 selects between these binding targets is crucial for elucidating its immune evasion strategy. Therefore, accurate estimation of binding parameters is essential for gaining mechanistic insights into these interactions. However, we observe heterogeneous binding in the BLI analysis of the NS1-TRIM25 interaction.

By combining BLI with isothermal titration calorimetry (ITC) and size-exclusion chromatography coupled with multi-angle light scattering (SEC-MALS), we characterize the nature of heterogeneous binding and its impact on the analysis of the NS1-TRIM25 interaction. Furthermore, we propose an alternative approach to mitigate the adverse effects of heterogeneous binding, improving the accuracy of binding parameter estimation.

## Results and Discussions

### Characterization of heterogeneous binding

To monitor the binding between IAV NS1 and TRIM25, we immobilized NS1 as a ligand on the biosensor and dipped it into a buffer containing TRIM25 as the analyte. Overall, the NS1-TRIM25 interaction was best described as a biphasic process, consisting of a rapid primary binding event followed by a slower secondary binding process, i.e., heterogeneous binding during which the BLI signal continuously drifted upward instead of reaching a steady-state (**Fig 2A**). Surprisingly, this signal drift continued without approaching a plateau, even when the association period was extended to 2400 seconds (**Fig 2A**).

**Figure 2:**
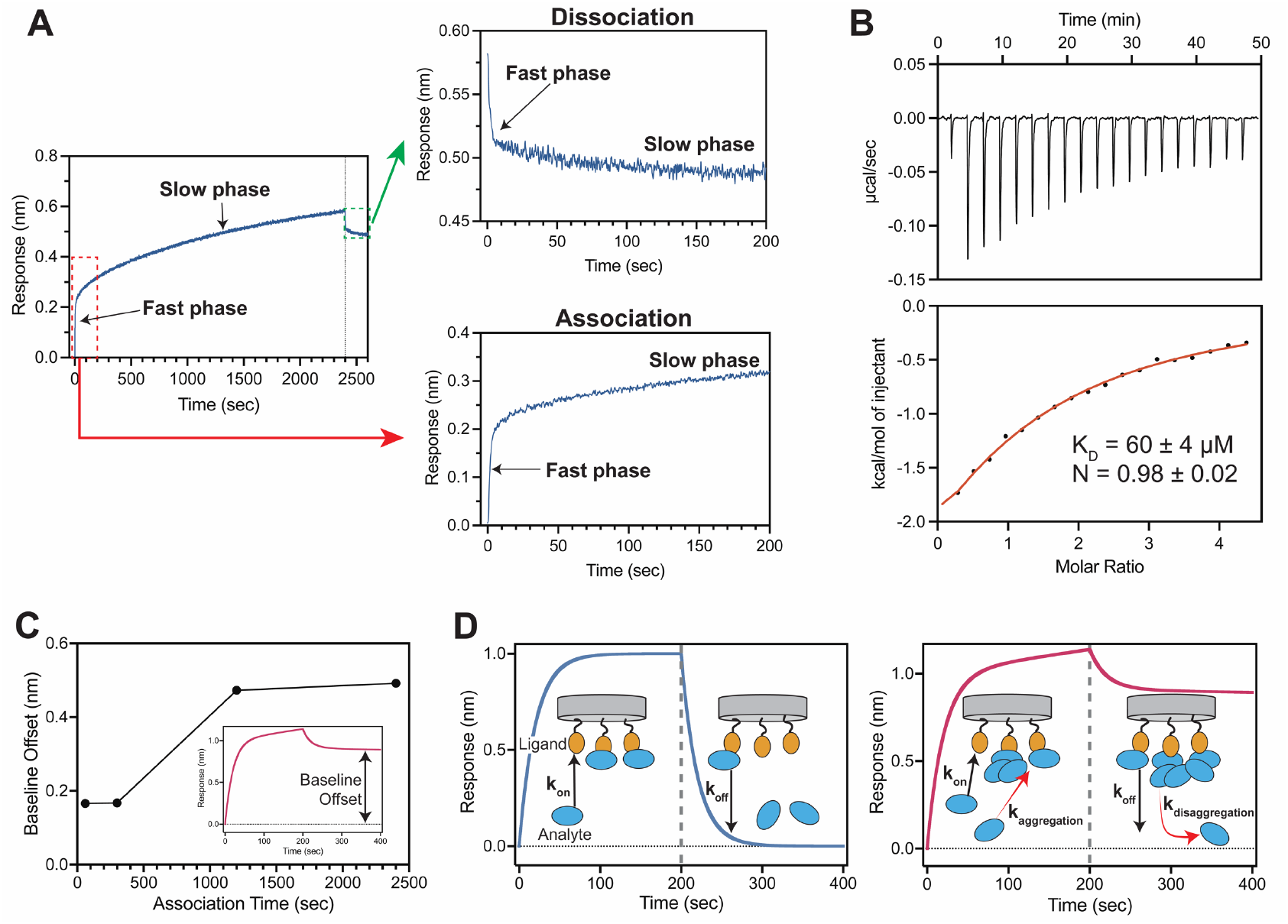
Effects of heterogeneous binding. **(A)** BLI sensorgram of TRIM25 binding to NS1. The first 200s of association and dissociation are shown in separate graphs. (**B)** ITC thermogram and isotherm for the interaction between NS1 and TRIM25. The red line indicates the fit curve for a 1:1 binding model. Numbers after ± symbol represent the standard deviation of two measurements. **(C)** Measurement of baseline offset as a function of association time. Definition of baseline offset is shown in inset on representative sensorgram. **(D)** A schematic showing ideal (left panel) and heterogenous (right panel) binding. Dotted vertical line demarcates beginning of dissociation phase. Analytes aggregate with pre-formed analyte-ligand complexes during association phase. These aggregated complexes dissociate slowly resulting in baseline offset (right panel).

To understand the basis of the heterogeneous binding, we first examined whether NS1 and TRIM25 form a functional multimeric complex, which may exhibit two distinct binding kinetics. The crystal structure of the NS1-TRIM25 complex revealed that TRIM25, as a dimer, binds to two NS1 molecules, i.e., a 1:1 binding stoichiometry (**Fig 1A**).^27^ To further characterize the NS1-TRIM25 interaction, we employed isothermal titration calorimetry (ITC), a gold-standard method for determining binding stoichiometry. The ITC results showed a binding stoichiometry (N) close to 1 (N = 0.98 ± 0.02), indicating the absence of higher-order NS1-TRIM25 interactions in solution (**Fig 2B**).

We next tested whether the observed signal drift was caused by nonspecific binding (NSB) of free analytes to the biosensor surface. NSB-induced signal changes are particularly concerning when analyzing weak protein-protein interactions (PPIs), as such studies often require high analyte concentrations. However, NSB by TRIM25 was minimal due to the presence of an NSB blocker (**Supplementary Fig 1**).^10^

Another possible explanation for the observed heterogeneous binding behavior could be the surface binding of oligomeric analytes. In other words, in addition to fast 1:1 binding, some analytes may slowly form oligomers and subsequently bind to a ligand. To assess whether TRIM25 or NS1 alone form higher-order multimers that could contribute to heterogeneous binding, we performed size-exclusion chromatography coupled with multi-angle light scattering (SEC-MALS). Although we incubated the mixture of NS1 and TRIM25 for 2 hours before applying it to SEC-MALS, the result revealed only two peaks corresponding to the expected TRIM25 dimer and NS1 monomer (**Supplementary Fig 2**). These results indicated that both TRIM25 and NS1 are homogeneous in free solution, excluding heterologous oligomerization as a source of heterogeneous binding.

We also observed that heterogeneous binding prevented the dissociation curves from returning to baseline (i.e., a BLI signal of 0 nm). Namely, the dissociation of heterogeneous binding complexes was extremely slow, suggesting that the ligand-analyte complexes remained practically irreversibly bound to the biosensors (**Fig 2A**). Furthermore, due to incomplete dissociation, the baseline offset from 0 nm increased with the length of the association period, indicating a time-dependent aggregation process during the association phase (**Fig 2C**). We next examined whether the slow dissociation phase is the intrinsic feature of the NS1-TRIM25 complex. If the dissociation of the complex is also slow in solution, the resulting protein complexes would be detectable by SEC-MALS. However, we did not observe such complexes when a mixture of NS1 and TRIM25 was analyzed, even after incubating the mixture for 2 hours (**Supplementary Fig 2**). This finding suggests that the functional NS1-TRIM25 complex dissociates rapidly in solution, whereas the slow dissociation phase observed in the BLI data likely reflects the desorption of surface-induced protein aggregates. Moreover, this observation suggests that monitoring changes in the dissociation baseline as a function of the association period could be a simple diagnostic test for detecting aggregation.

Taken together, our results indicated that heterogeneous binding is most likely due to the slow aggregation of excess analytes around pre-formed analyte-ligand complexes (**Fig 2D**). Adsorption of these heterogeneous protein aggregates onto the biosensor surface may lead to near-irreversible binding, as reflected in the extremely slow dissociation phase (**Fig 2A**), which was not observed in solution using ITC and SEC-MALS. This finding is somewhat unexpected, as it is commonly assumed that the inclusion of BSA or detergents (e.g., Tween 20) in the buffer would prevent aggregation.^32, 33^ Our results demonstrate that this assumption may not hold true for certain proteins.

### Effects of heterogeneous binding on kinetic analysis

Although BLI provides both steady-state and kinetic approaches for estimating K_D_, the steady-state approach is more challenging for weak PPIs. Accurate K_D_ determination via the steady-state approach requires a broad range of analyte concentrations, typically spanning 0.1 × K_D_ to 10 × K_D_.^3^ Given the limited solubility of many proteins, achieving this condition can be difficult. Moreover, NSB becomes more severe as the analyte concentration increases.^10^ Therefore, the kinetic approach is generally preferred for characterizing weak PPIs. However, our study indicated that heterogeneous binding disrupts both association and dissociation signals (**Fig 2A**). These anomalous binding curves may lead to erroneous data fitting unless the effects of heterogeneous binding are properly accounted for. Similar effects on the dissociation curves from varying association phase times were also reported in SPR studies.^11, 12, 34^

Therefore, we tested the impact of heterogeneous binding on the analysis of association kinetics. We first noticed that the k_obs_ value obtained using a single exponential function varied significantly with the length of the association period, differing by around 100-fold between 60 and 2400 seconds (**Fig 3A**). In contrast, fitting with a biphasic function resulted in only about 3-fold difference over the same time range. This result indicated that a biphasic function could result in accurate fitting when heterogenous binding is present in the binding data. We further tested whether a biphasic function accurately separates the kinetic parameters of the NS1-TRIM25 interaction from the slow heterogeneous binding. Indeed, the linear fit of the plot of k_obs_ vs [analyte] resulted in k_on_ and k_off_ values of 0.006 ± 0.001 s^-1^µM^-1^ and 0.37 ± 0.03 s^-1^, respectively (**Fig 3B**). The resulting kinetic K_D_ value (K_D_ = k_off_ / k_on_) was estimated to be 62 ± 11 μM, which is in excellent agreement with the ITC result of 60 ± 4 µM (**Fig 2B**).

**Figure 3:**
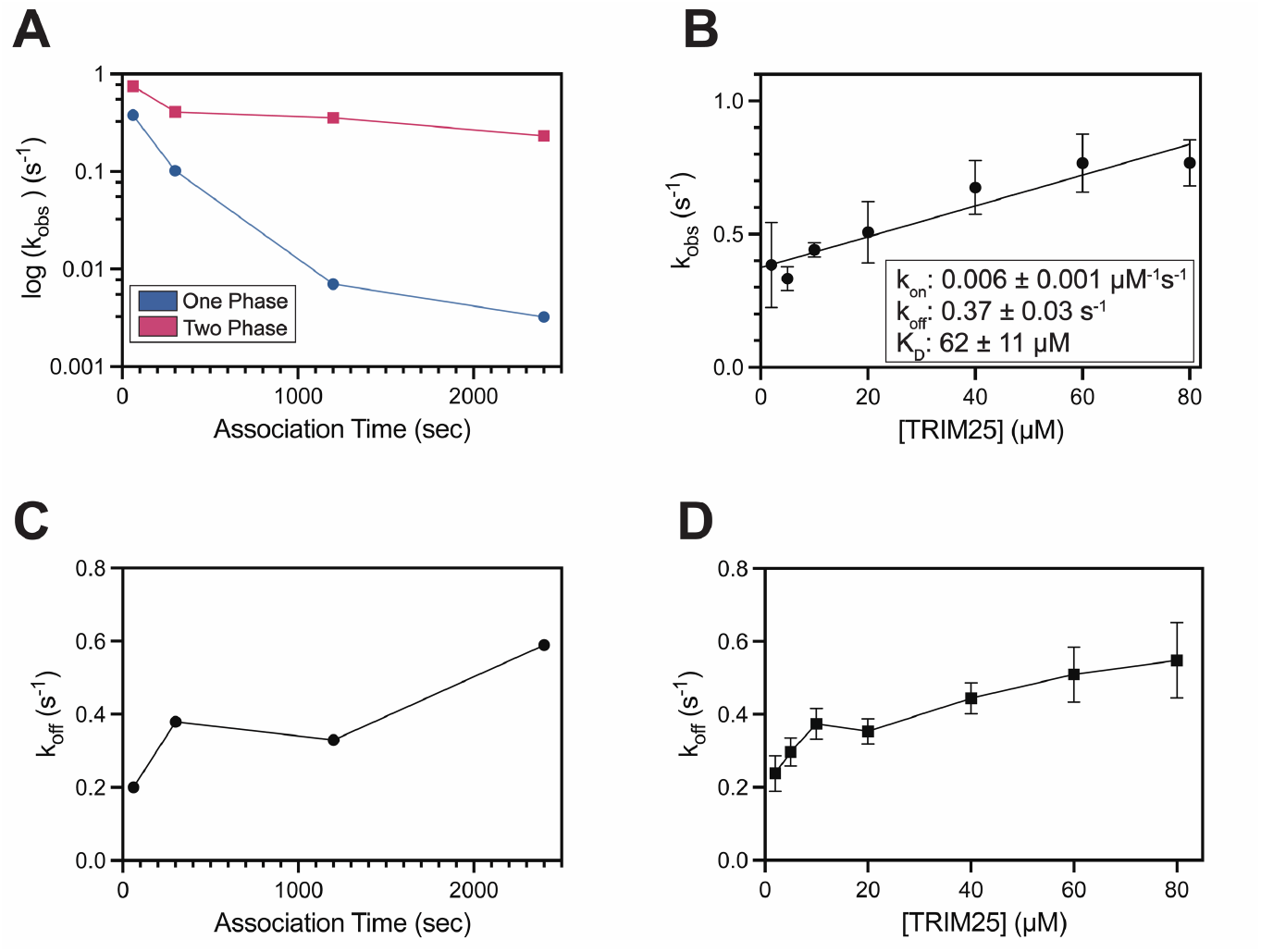
Heterogenous binding affects kinetic analysis. **(A)** k_obs_ values as a function of the association period. Blue and magenta symbols represent the k_obs_ values obtained from single- and biphasic-exponential fits, respectively. **(B)** Plot of k_obs_ vs TRIM25 concentration. Error bars represent the standard deviation from three repeated measurements. Numbers after ± symbol represent the standard deviation of three repeated measurements. **(C-D)** k_off_ values as a function of **(C)** association time and **(D)** analyte concentration. k_off_ is calculated from the fast phase of a biphasic exponential fit. Error bars represent the standard deviation from three repeated measurements.

We next examined the effects of heterogeneous binding on the analysis of dissociation kinetics. Direct measurement of k_off_ from dissociation curves is critical, as the k_off_ value obtained from a linear fit of k_obs_ vs. [analyte] can be unreliable due to errors associated with extrapolating the fit curve to the y-axis.^35^ We noted that heterogeneous binding resulted in incomplete dissociation, necessitating the use of an artificial baseline in the fitting process (**Fig 2A** and **2D**). Moreover, the extent of heterogeneous binding affected the fitting results even when a biphasic function was applied. For example, the k_off_ values increased threefold (0.2 s^−1^ to 0.6 s^−1^) between the shortest (60 s) and longest (2400 s) association periods (**Fig 3C**). We also observed that k_off_ values increased as a function of analyte concentration (**Fig 3D**). Based on these results, we recommend measuring k_off_ at the lowest possible analyte concentration and with the shortest possible association period to minimize the influence of heterogeneous binding, provided that the BLI signal change remains sufficient for quantitative analysis.

The analysis of dissociation process is often considered straightforward, based on the assumption that heterogeneous binding only affects the association phase. Contrary to this common perception, we found that heterogeneous binding also significantly affects the analysis of dissociation curves. This is important because dissociation analysis is critical for drug discovery, as residence time is often an indicator of a robust pharmacological response.^36, 37^

### Alternative Approach to Reduce the Adverse Effects of Adsorption

When kinetic parameters are estimated by a linear fit of k_obs_ vs [analyte], a broad range of [analyte] improves the accuracy of the fitted parameters. However, when heterogeneous binding is severe, this approach can be challenging. For example, since heterogeneous binding is more pronounced at higher analyte concentrations (**Fig 3D**), extrapolating the concentration-dependent linear fit might lead to erroneous k_off_ values.

Therefore, as an alternative, we tested whether kinetic parameters could be reliably determined using a single analyte concentration where heterogeneous binding is minimal. Estimating kinetic parameters from a single analyte concentration is often associated with high uncertainty. To address this, we repeated the measurements multiple times, providing the same degree of freedom (df = 6) as the concentration-dependent approach.

We first tested 20 μM TRIM25, which is lower than the K_D_ value (60 μM) and exhibited mild heterogeneous binding while still producing a reasonable BLI signal amplitude upon binding to NS1. The single concentration approach yielded 0.007 ± 0.004 μM^−1^ s^−1^ and 0.33 ± 0.07 s^−1^ for k_on_ and k_off_ values, respectively (**Fig 4A**). To further validate our approach, we tested the approach using 10 μM TRIM25, which is significantly lower than the K_D_ value (**Fig 4B**). Remarkably, the fitting result (k_on_ = 0.008 ± 0.006 s^-1^ µM^-1^ and k_off_ = 0.29 ± 0.02 s^-1^) was virtually identical to that at 20 μM TRIM25. These results were also in excellent agreement with values obtained from concentration-dependent measurements and ITC (**Fig 3B**). Consequently, these results suggest that kinetic measurements using a single analyte concentration can serve as a viable alternative when heterogeneous binding interferes with standard concentration-dependent analysis. It should be noted, however, that a concentration-dependent approach generally provides more accurate kinetic parameters than using a single analyte concentration. That said, when properly executed, a single concentration approach can be an alternative in cases of heterogeneous binding.

**Figure 4:**
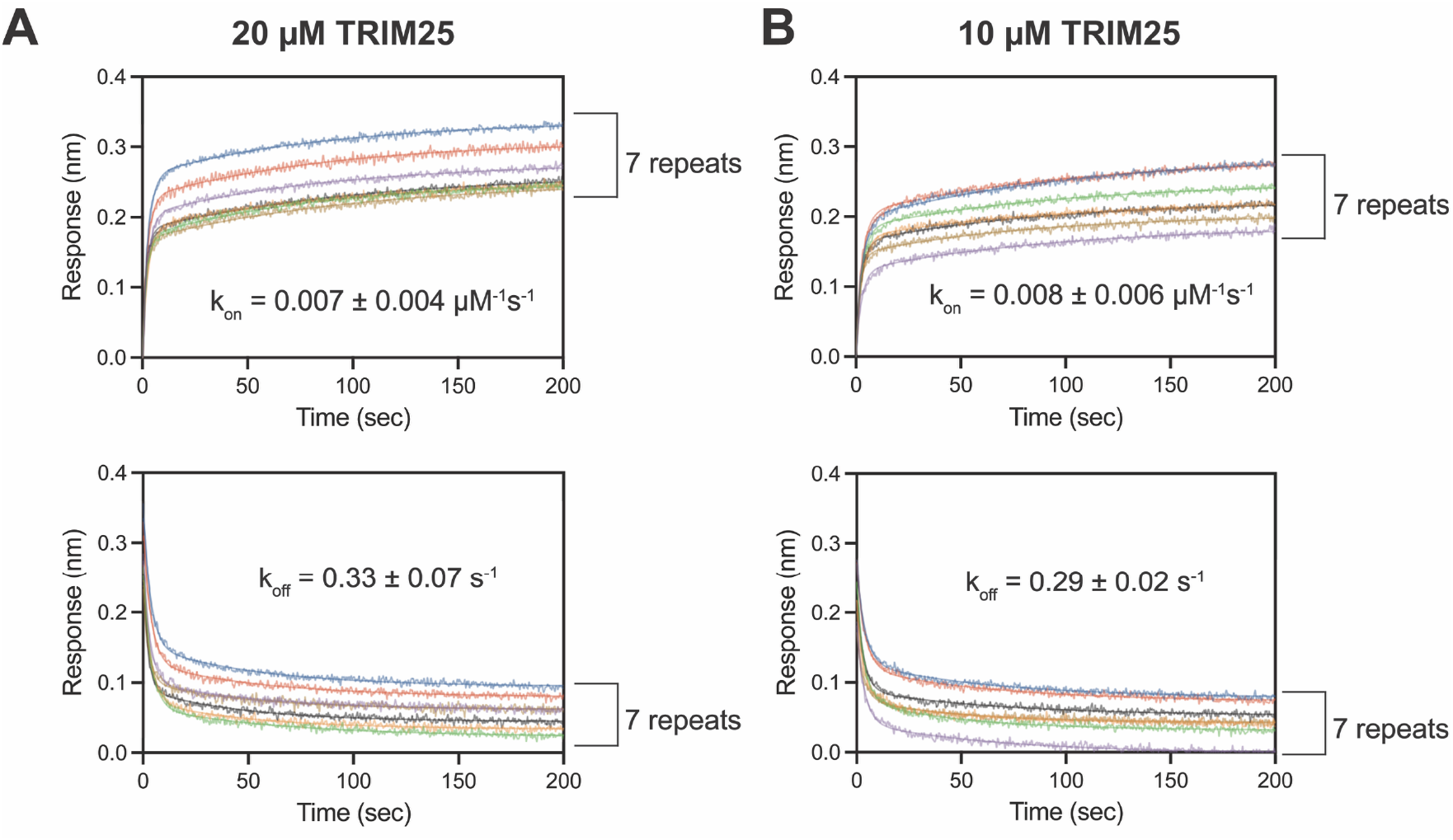
BLI at a single analyte concentration. BLI sensorgrams for **(A)** 20 µM TRIM25 and **(B)** 10 µM TRIM25. Seven repeated measurements are shown in different colors. Numbers after ± symbol represent the standard deviation of seven repeated measurements. Data is fitted with a biphasic exponential equation.

## Conclusions

It is often assumed that the analyte concentration should range from 0.1 × K_D_ to 10 × K_D_, even for kinetic experiments.^38^ While this condition is critical for steady-state K_D_ measurements and desirable for kinetic K_D_ as well, artifacts arising from heterogeneous binding often make it challenging to achieve these conditions. This issue is further exacerbated when characterizing weak PPIs, which requires high analyte concentrations to satisfy the conditions.

Our study indicated that non-specific aggregation of extra analytes with a ligand-analyte complex on the surface of biosensors can cause heterogeneous binding (**Fig 2D**). Significantly, heterogeneous binding adversely affects both the association and dissociation processes (**Fig 2A**). Moreover, we found that heterogeneous binding increases with the association period and analyte concentration. Accordingly, we proposed measuring the offset of dissociation baseline as a function of association time and analyte concentrations to test for the presence of heterogeneous binding. This simple test will provide justification for using a biphasic equation for the data analysis.

We also demonstrated that kinetic binding parameters (k_on_, k_off_, and K_D_) can be reliably measured using repeated measurements at a single analyte concentration, where heterogeneous binding is minimized. In favorable cases, reliable binding parameters could be obtained at concentrations significantly lower than K_D_.

## Materials and Methods

### Protein Expression and Purification

Genes encoding TRIM25 (residues 183-380) and the Puerto Rico 8 (PR8) IAV NS1 (residues 80-205) were prepared by the gene-synthesis service from Genscript. The NS1 protein contains a W187A mutation to help prevent protein aggregation.^39^ All proteins were expressed with an N-terminal His_6_ and SUMO tag in BL21 (DE3) *E. coli* cells. In addition, NS1 contained an Avi tag, and was subsequently co-transformed with the BirA plasmid for biotinylation during expression. All proteins were purified as described previously.^10, 31^ Briefly, TRIM25 was purified via a His-Trap HP Ni NTA column (Cytiva), followed by SUMO protease to cleave the tag, and a final His-Trap HP Ni NTA column before dialysis and storage. NS1 was purified in a similar way, with the additional step of gel filtration chromatography using a HiLoad 16/600 Superdex 200 pg column (GE Healthcare). For biotinylated protein, gel filtration chromatography was substituted in lieu of SUMO protease cleavage. Protein purity was confirmed by SDS-PAGE, with all samples >95% pure. All reported concentrations are given as monomer concentrations to reflect the 1:1 binding stoichiometry of TRIM25 to NS1.

### Isothermal Titration Calorimetry

All ITC experiments were recorded at 25 ºC using a Microcal-PEAQ-ITC calorimeter (Malvern Panalytical). Experiments were carried out in buffer containing 20 mM sodium phosphate (pH 7), 100 mM NaCl, and 2 mM TCEP. TRIM25 (500 µM) was injected into NS1 (25 µM) in a series of 18, 2 µL injections. All data was fit using a 1:1 binding mode to obtain the reported thermodynamic parameters using the software provided by the instrument. Reported parameters are the average and standard deviation of two independent measurements.

### Size Exclusion Chromatography with Multi-Angle Light Scattering (SEC-MALS)

20 µM TRIM25 was incubated with 160 µM of NS1 for 2 hours at 25 ºC. The protein complex was then injected into a Superdex 75 Increase 10/300 GL column (GE Healthcare) connected to a miniDAWN MALS and an Optilab dRI detector (Wyatt Technology) using a running buffer of 20 mM sodium phosphate (pH 7), 100 mM NaCl, and 2 mM TCEP. The molecular weight was determined through the ASTRA software package v8.2.2 (Wyatt Technology).

### Biolayer Interferometry

The binding of surface-immobilized NS1 to TRIM25 was measured using a Gator Pilot biolayer interferometry instrument (Gator Bio) at 25 ºC. Biotinylated NS1 was immobilized onto streptavidin biosensor probes, and experiments were conducted in buffer containing 20 mM sodium phosphate (pH 7), 150 mM NaCl, 1% BSA, 0.6 M sucrose, and 1 mM TCEP. All experiments were conducted at 1500 rpm with a 10 Hz acquisition rate. Tips were regenerated with a solution of 0.5 M guanidinium chloride and 0.6 M sucrose to remove bound TRIM25 and protein aggregate from immobilized NS1. Regeneration efficiency was > 98% effective. Error is given as the standard deviation of 3 runs for the multi-concentration data and 7 runs for the single-concentration data. All BLI data were analyzed using GraphPad Prism 10 (GraphPad Software). For the single concentration data, k_off_ is first calculated from dissociation and then used to obtain k_on_ from association (i.e.,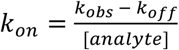). Equations 1 and 2 were used for single and biphasic fits to the data, respectively:

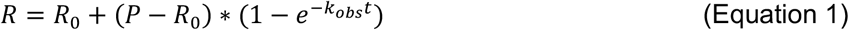

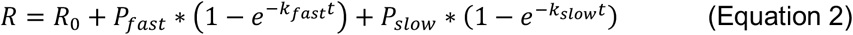

where R_0_ represents the response value at time = 0, P represents the response value at saturation, k is the observed rate constant, and t is time. Both P and k can be further classified into “fast” or “slow” in a biphasic fit. Equations 1 and 2 are represented in GraphPad Prism 10 as “one phase” and “two phase” association/dissociation respectively.

## Acknowledgments

We thank Meega Reji for her assistance in purifying NS1.This work was funded by NIH grant R53GM152007 and the Welch Foundation grant A-2028-20230405.

