## Supplementary Figures for "Overcoming effects of heterogeneous binding on BLI analysis"

Supplementary Figures 1 and 2.

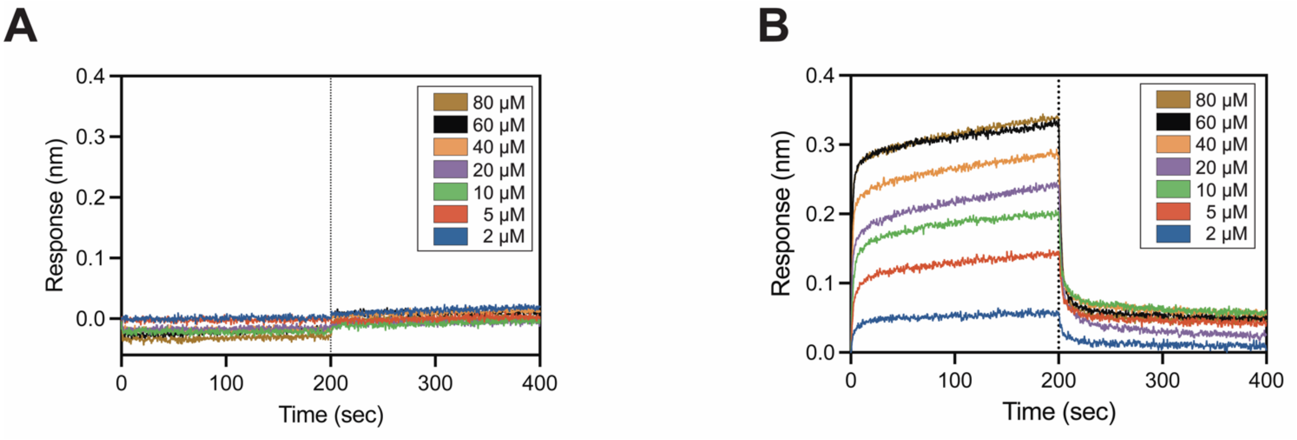


**Supplementary Figure 1. (A)** BLI Sensorgram of non-specific binding (NSB) at various concentrations of TRIM25. NSB is the BLI response in the absence of immobilized ligand. **(B)** Representative BLI sensorgram of the binding between NS1 and TRIM25. The dotted line demarcates the initiation of the dissociation phase.

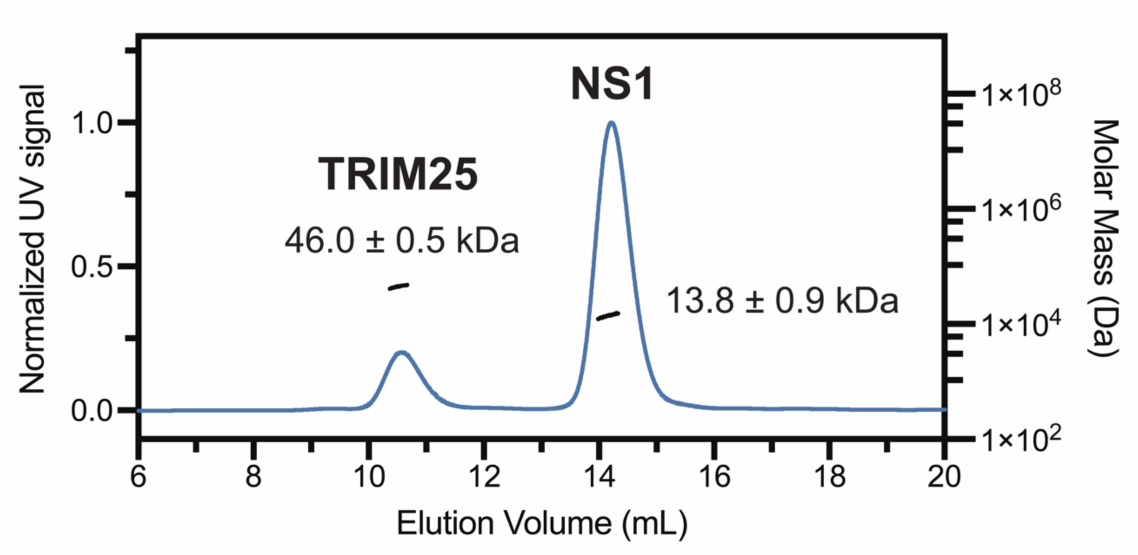


**Supplementary Figure 2.** SEC-MALS chromatogram of the interaction between TRIM25 and NS1. The theoretical molecular weights of the NS1 (monomer) and TRIM25 (dimer) constructs are 13.03 and 45.54 kDa respectively. No complex or aggregates were observed after a two-hour incubation.
